# Peripheral B Lymphocyte Serves as A Reservoir for The Persistently Covert Infection of Mandarinfish *Siniperca chuatsi* Ranavirus

**DOI:** 10.1101/2024.05.06.592682

**Authors:** Wenfeng Zhang, Hui Gong, Qianqian Sun, Yuting Fu, Xiaosi Wu, Hengwei Deng, Shaoping Weng, Jianguo He, Chuanfu Dong

## Abstract

Genus *Ranavirus* in family *Iridoviridae* is composed of large members with various genomic sizes and viral gene contents, infecting a variety of ectothermic vertebrates including reptile, amphibians and bony fish worldwide. Mandarinfish ranavirus (MRV) is a very distinctive member among genus *Ranavirus*. Persistently convert infection of MRV were previously observed in natural outbreak of MRV, but the underlying mechanism remains unclear. We here evidenced that mandarinfish peripheral B lymphocytes are implemented as viral reservoirs to maintain persistent and covert infection. When mandarinfish were infected with sublethal dosage of MRV under nonpermissive temperature (19 ºC) and permissive temperature (26 ºC), respectively, all fish in 19 ºC group survived and entered persistent phase of infection characterized with very low viral load in white blood cell, whereas partial fish died of MRV infection in 26 ºC group, and the survivals then initiated persistent status. Gradually raising temperature, vaccination and dexamethasone treatment can reactivate the quiescent MRV to replicate and result in partial mortality. The viral reservoir investigates showed that IgM^+^-labelled B lymphocytes but not CD3Δ^+^-labelled T lymphocytes and MRC-1^+^-labelled macrophages are target cells for the persistent infection of MRV. Moreover, the quiescent MRV could not be reactivated by heat-killed *Escherichia coli*, indicating a very different reactivation mechanism from that of other known rannaviral member. Collectively, we are the first to confirm the presence of B cell-dependent persistent and covert infection of ranavirus, and provide a new clue for better understanding the complex infection mechanism of vertebrate iridovirus, especially regarding ranavirus.

**IMPORTANCE:** Viruses known as HIV, HBV and EBV etc. evade host immune clearance by establishing long-term even lifelong persistent or latent infection. In vertebrate iridovirus, FV3, the type species of genus *Ranavirus* was evidenced to establish persistent infection by using *Xenopus* peritoneal macrophages as reservoirs. MRV is a very distinctive ranavirus from FV3 with very different genomic content and host species. We here uncovered MRV establishes persistent and covert infection by using peripheral B lymphocytes as virus reservoirs. During persistent infection, very low copies of quiescent MRV were harbored in peripheral B lymphocytes. Water temperature stress, vaccination stimulation, and dexamethasone treatment can reactivate quiescent MRV to replicate in abundance via a non-TLR5-mediated manner, and results in recurrence of MRV disease. Our finding suggests the diversity and complexity of the pathogenic mechanisms among ranaviruses, and also has important scientific significance for in-depth understanding of the infection and immunity interaction of vertebrate iridoviruses.

## INTRODUCTION

Iridovirus is a large viral family, with a linear double-stranded DNA (dsDNA), genome sizes ranging from 98 kb to 220 kb and encoding open reading frames (ORFs) ranging from 98 to 221(1-4). The *Iridoviridae* family can be divided into two subfamilies, namely, *Alphairidovirinae* and *Betairidovirinae*, and also are referred to vertebrate iridovirus (VI) and invertebrate iridovirus (IVI), respectively(2). Among *Alphairidovirinae*, the *Ranavirus* genus has the most abundant members, with great diversity in genome content, the most diverse array of hosts, including bony fish, reptiles and amphibians, and very complex pathogenesis mechanisms(5-7). Ranavirus-associated viral diseases have caused considerable economic loss and ecological disasters worldwide due to their high contagiousness and lethality to various economically valuable bony fish species(6-8).

Mandarinfish ranavirus (MRV), also known as a variant of largemouth bass virus (LMBV) or Santee-Cooper ranavirus (SCRaV), has being emerged as an important pathogenic threat to several bony fish species. The known sensitive natural hosts of MRV/LMBV include largemouth bass (*Micropterus salmoides*)(9, 10), koi carp (*Cyprinus carpio*)(11), barcoo grunter (*Scortum barcoo*)(12), two tropical ornamental fish, Doctor fish (*Labroides dimidatus*) and Guppy (*Poecilia reticulata*)(13) as well as mandarinfish (*Siniperca chuatsi*)(3, 14, 15). In mandarinfish, MRV infection generally manifests in two distinctive clinical phenotypes: acute and chronic(4, 14, 15). During acute infection, morbidity generally occurs at 2-4 days post-infection, and is characterized by vomiting and then severe ascites syndrome(15). Subsequently, mass mortalities ensue within 3-10 days with an accumulative mortality of >50%(14, 15). The tropism tissues during acute infection includes pyloric caeca, spleen, kidney, liver and intestines(4, 14). In contrast, the chronic phase of infection was featured by lethargy, anorexia, abnormally swimming and impaired coordination, and could persist for up to 5 months(14). During chronic infection, the most surviving fish become emaciated and deformed due to long-term anorexia or abnormal feeding, and then lose their aquaculture value. Especially, during chronic infection, very low copes of genomic DNA were detected just by real-time quantitative PCR but not by conventional PCR using the visceral organs such as spleen, liver and kidney as target tissues. We previously designated this long-lasting, undetectable status of MRV-carrier by conventional PCR detection as recessive carrier(14) and currently designated it as persistently covert infection (PCI). During PCI, sporadic death but not mass mortality became the major events.

Prior to the emerging MRV, infectious spleen and kidney necrosis virus (ISKNV), the type species in genus *Megalocytivirus of Alphairidovirinae*, has been considered as the first important pathogenic threat to mandarinfish industry(16). Until recently, the inactivated ISKNV vaccine with an absolute protection effect of >90% was developed and officially licensed for a commercial application(14, 17). The inactivated ISKNV vaccine also showed the same high efficacy in Asia seabass *Lates calcarifer* and spotted seabass *L. maculatus* against various genotypes of ISKNV/RSIV isolates in ISKNV species(18, 19). Unexpectedly, we attempted to develop an effective MFF-1 cell-based MRV vaccine conducted along with the same rational technology routine as that of the inactivated ISKNV vaccine, but failed. Almost no protection was achieved by the inactivated MRV vaccine, regardless of MRV antigen dosage (unpublished data by Dong’s lab). Generally, ISKNV/RSIV is proposed to be a transient pathogen. As a result, either the inactivated vaccine or non-lethal active virus exposure could elicit robust immune protection against the infection or re-infection with virulent ISKNV/RSIV(17, 20-22). Conversely, MRV displays to have characteristics of persistent infection. As an opposite result, the inactivated MRV vaccine confers no protection. Additionally, the non-lethal active MRV exposure could cause long-term PCI(14).

White blood cell (WBC), a pivotal role in the immune system, serves as dormant or latent carriers for numerous viruses. For instance, the human immunodeficiency virus (HIV) specifically targets T cells as its latent host cells, while the human T-cell leukemia virus (HTLV) can establish residence in both B and T cells(23, 24). Similarly, the herpesvirus can conceal itself within B lymphocytes in both human and animal(25-29). These viruses strategically target the immune system for infection and latency, enabling them to elude immune clearance and persist within the host for extended periods. Studies have underscored the significance of macrophages as persistent hosts for amphibian ranaviruses(30-33). FV3 is believed to undergo quiescent infection, evident in the detection of FV3 DNA within peritoneal leukocytes following initial infection(34, 35). During quiescent state, the inactive viral particles are only identified within peritoneal macrophages(36). The injection of heat-inactivated *E. coli* can reactivate FV3 in asymptomatic adult frogs and lead to fatal infections(37, 38). Besides FV3, no other ranavirus has been studied comprehensively on its persistent infection and reactivation. As a very distinctive rannaviral member, we here revealed that MRV has a unique characteristic of B cell-targeted persistent infection, which is not only completely different from that of the well-studied megalocytiviral ISKNV/RSIV but also quite different from that of ranaviral FV3.

## MATERIALS AND METHODS

### Animals, virus and antibodies

Three-month old mandarinfish with body weight of 200 g∼300 g per fish were obtained from a fish farm in Foshan, Guangdong Province, China. All animal experiments were conducted in accordance with the animal testing procedures of Guangdong Province, China, and were approved by the Ethics Committee of Sun Yat-sen University (Approval number: SYSU-LS-IACUC-2024-0016). Before experiment, mandarinfish were randomly sampled and tested to confirm free of asymptomatic MRV carrier by qPCR. MRV isolates ZQ17 were isolated and characterized from a mass mortality event of mandarinfish(14). The mandarinfish fry (MFF-1) cell line was grown in complete Dulbecco’s modified Engle’s medium (DMEM) (Gibco) containing 10% fetal bovine serum (FBS) (Gibco) at 26°C containing 5% CO2 to amplify MRV(39). Anti-mouse monoclonal antibody (mAb) of MRV 1C4 was constructed in our recent report(4). Anti-mouse mAb of mandarinfish IgM 7F12F6 was kept in our lab(40). Polyclonal antibodies (pAb), specific to the paired domain of mandarinfish Pax5, specific to mandarinfish T-cell surface glycoprotein CD3 delta chain and specific to mandarinfish mannose receptor C-type 1 were prepared and stored in our laboratory (data not shown).

### Microchip label and artificial infection

Mandarinfish were divided into two groups (**Fig. 1A**). Group I was comprised of 15 fish, and reared in an aquatic environment with a constant water temperature of 26°C, while group II consisted of 20 fish and was maintained in a water temperature environment of 19°C. Prior to infection, individual mandarinfish in group II were labeled M1∼M20 by injecting microchip into back muscle. Following two weeks of acclimatization, all fish were challenged by intraperitoneal injection with MRV-ZQ17, 10^6.5^ TCID_50_/0.1 mL/fish. Daily monitoring of morbidity and mortality was conducted over the subsequent months. Fish were fed with a daily diet of live commercial bait fish.

**FIG. 1.**
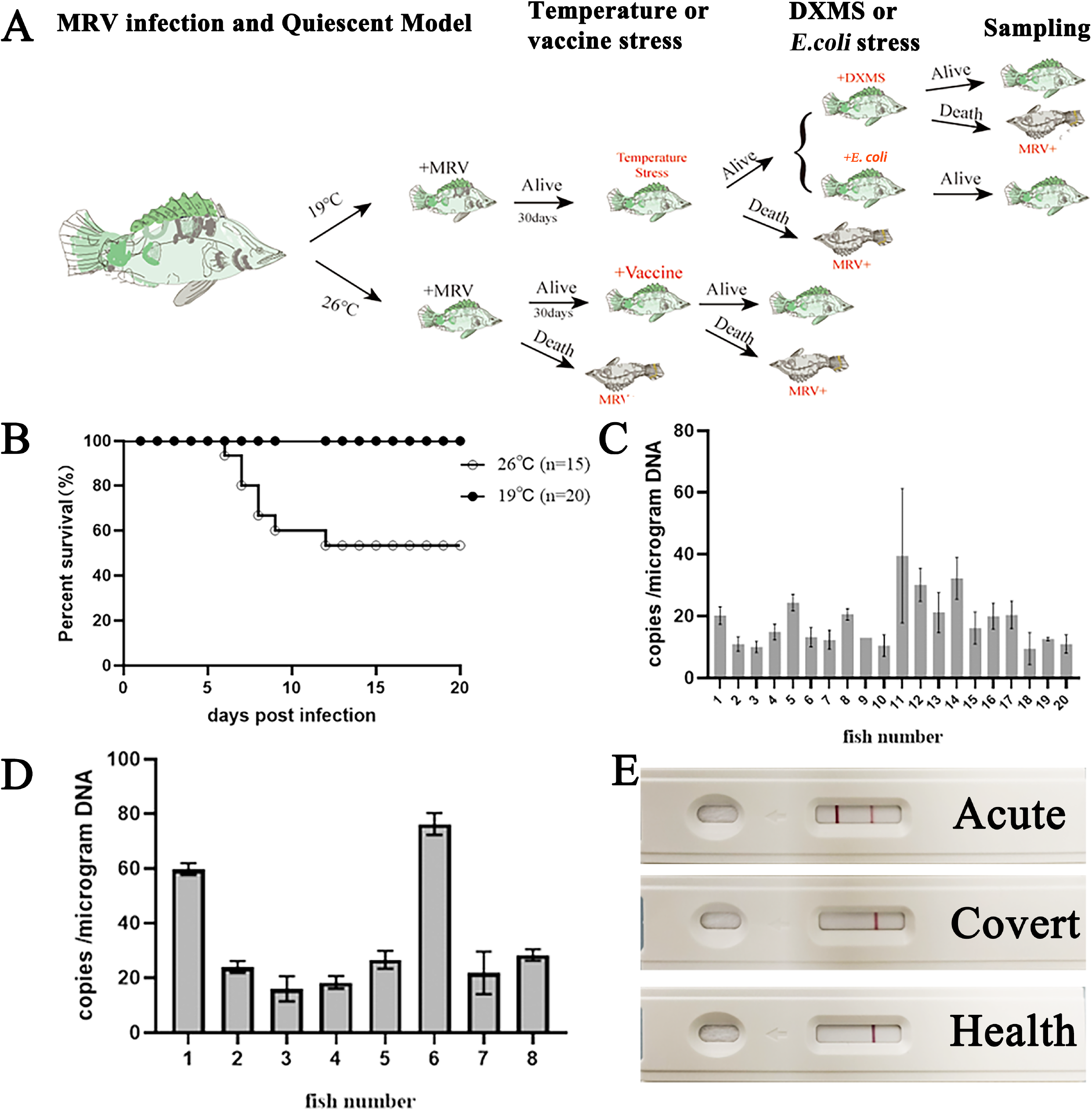
Tracking the status of MRV-infected mandarinfish under different temperature conditions. (A) The experimental outline of MRV infection and quiescent model. Briefly, mandarinfish was infected under two different temperature conditions. In 19 °C infection group, all fish survived and receive raising temperature stress at 30 dpi. Dead fish was confirmed to die of MRV infection. After acclimatization, the survivals were treated with dexamethasone (DXMS) and killed *E. coil*, respectively. In 26 °C infection group, partial mortality occurred with 12 dpi. The survivals were received vaccination with the inactivated MRV vaccine. Dead fish was confirmed to die of MRV infection. (B) The survival curve of MRV infected mandarinfish under 19°C and 26°C. All fish survival in 19°C infection group, and partial mortality was observed in 26°C in infection group. (C) and (D) The MRV DNA copy number per microgram of total WBC DNA isolated from 20 mandarinfish under 19°C group and 8 survival fish under 26°C at 30 dpi, respectively. (E) Detection of MRV using anti-MRV mAb-based colloidal gold-immunochromatographic fast detection strip. The positive MRV was obtained just from the dead fish suffering the primary infection of MRV in 26 °C infection group (Acute), but negative for 19 °C infection group and mock infection.

### Raising temperature stress

To determine whether the quiescent MRV can be reactivated by raising temperature stress, at 38 days post infection (dpi), water temperature of group II was incrementally increased from 19°C to 26°C with a gradual rate of 1°C per 2 days, ultimately maintaining a constant temperature of 26°C until the end of the experiment. (**Fig. 2A**). Subsequent to temperature elevation, daily observation was conducted to monitor pathological changes and mortality. The liver, spleen, kidney, and intestine from diseased fish were collected for quantitative PCR analysis to determine MRV load. Additionally, MFF-1 cells were inoculated to assess the development of cytopathic effects (CPE).

**FIG. 2.**
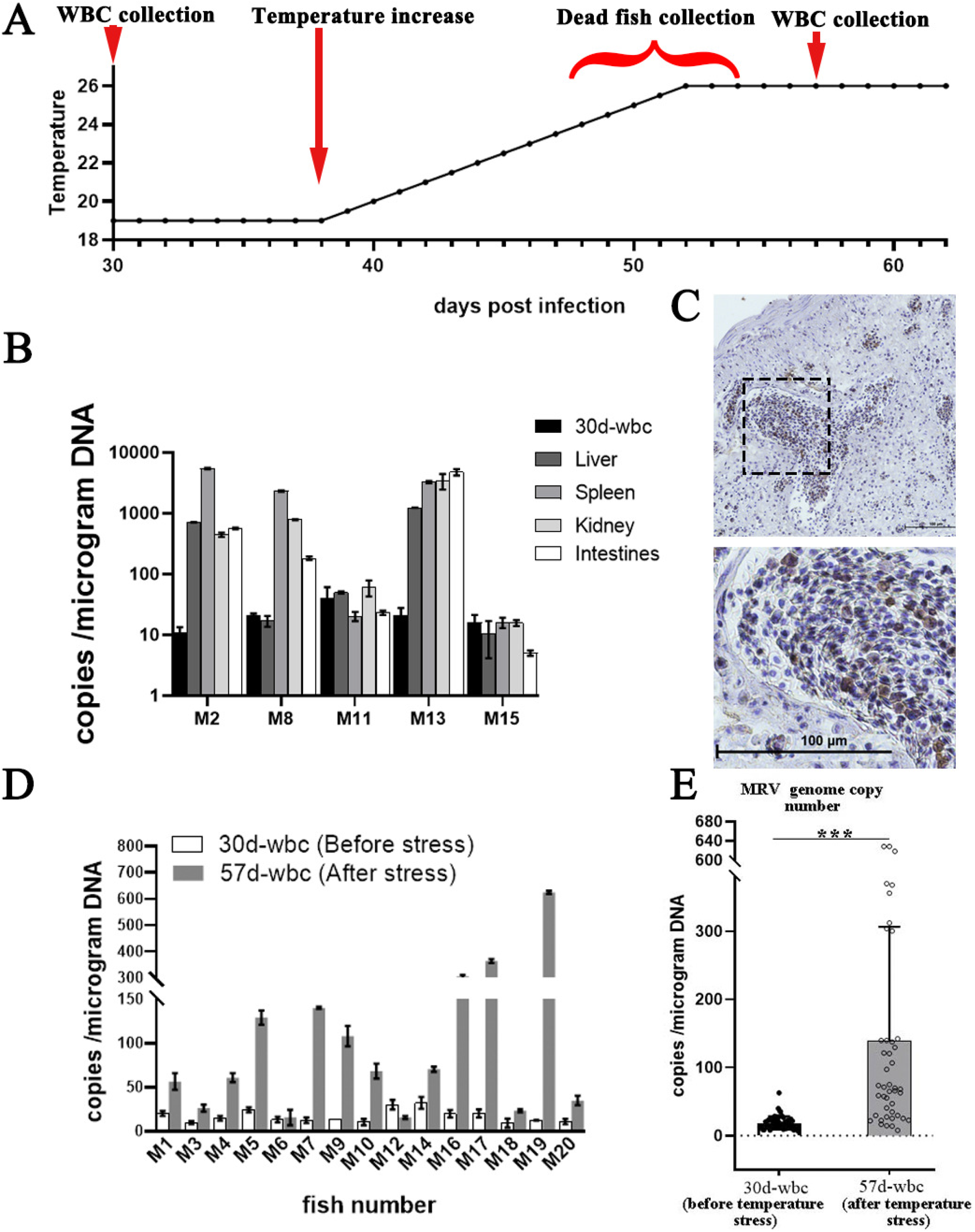
Reactivation of covert MRV via raising temperature stress. (A) The experimental outline of reactivation of covert MRV via raising temperature stress. The infected fish in 19 °C group were drawn blood for MRV load analysis at 30 dpi. After 7 days acclimatation, at 38 dpi, the water temperature was gradually elevated with 1 °C per two days. During raising temperature stress, dead fishes were sampled for MRV load measurement, and the survivals were conducted the second round of blood collection for MRV load assessment. (B) MRV DNA load in various tissues from 5 dead fish during the raising temperature stress. High MRV loads were observed in M2, M8 and M13. (C) Immuno-histochemistry of the intestines of dead fish M13. The tissue was recognized with mAb 1C4, and numerous infected cells were stained brown by DAB. Bar = 100 μm. (D) and (E), the MRV DNA copy number per microgram of total WBC DNA of each (D) and gross distribution (E), respectively. Fifteen survival mandarinfish on the 5th day after the completion of raising temperature stress and compared with that before stimulation. ***, P<0.001, in comparison with the average of MRV DNA before and after stimulation.

Concurrently, the fixed tissues were subjected for immunohistochemical analysis. When temperature has been sustained at 26°C for 5 days (at 57 dpi since the initial infection), blood samples were obtained from the surviving fish through tail vein following anesthesia for the purpose of WBC collection for MRV load detection (**Fig. 2A**).

### Vaccination, dexamethasone treatment and *E. coil* stimulation

MRV-ZQ17 isolate was used for vaccine preparation as previous description(41). Briefly, confluent MFF-1 cells were inoculated with MRV with a multiplicity of infection (MOI) of 0.1. When CPE was complete at about 3 dpi, whole cell suspensions were harvested and inactivated by formalin to a final concentration of 0.1% (v/v) at 4°C for 10 days. The inactivated virus was emulsified as water in oil (o/w) formation with injectable white oil (Esso, France). *E. coli* cells (DH5&alpha) were cultured overnight at 37°C, boiled for 1 h, pelleted by centrifugation, resuspended in PBS, and adjusted to a final concentration to 500,000 cells/mL. Finally, the PCI mandarinfish were injected by i.p. with 100 μL of MRV vaccine, 5.0 mg/kg of dexamethasone, or 100 μL of the resuspended heat-killed *E. coil*, respectively. During these treatments, tissues from moribund fish were collected for MRV load measurement.

### IgM^+^, CD3^+^ and MRC1^+^ WBC isolation and purification

WBC were collected after layering whole blood on a Ficoll-Paque Plus gradient according to the manufacturer’s instructions (GE Healthcare, United Kingdom) and washed twice with Hanks balanced salt solution (HBSS). Total WBC were stained firstly with anti-mandarinfish IgM, CD3 or MRC1 antibody on ice for 60 min and rinsed twice with HBSS. WBC were then stained with secondary Alexa Fluor 488-labeled goat anti mouse or rabbit IgG antibody with 1:500 dilution at 4°C for 60 min and washed twice, followed by positive and negative cells isolation using Beckman MoFlo Astrios EQs, according to manufacturer’s instructions (Beckman, America).

### Western blotting

The isolated WBC were lysed using RIPA lysis buffer (Pierce, China) and then subjected to 12% SDS-PAGE separation. The fractioned proteins were transferred for 90 min at 200 mA to nitrocellulose membranes (GE, USA). The mAb of mouse anti-mandarinfish IgM 7F12F6 and pAbs of rabbit anti-mandarinfish Pax5, CD3 and MRC1 were used as the first Abs. The horseradish peroxidase (HRP)-labeled-Goat anti-Mouse or Goat anti-Rabbit IgG (Sigma, USA) was used as secondary Ab. The recognized protein bands were visualized by 3,3 N-Diaminobenzidine tertrahydrochloride (DAB) solution staining (Roche, Germany).

### Confocal microscopy observation

The purified IgM^+^ cells and IgM^-^ cell residuals were fixed with 4% paraformaldehyde on tissue slides and permeabilized with 0.1% Triton X-100 in phosphate-buffered saline (PBS) containing 1% bovine serum albumin (BSA). The cells were stained with primary anti-IgM 7F12F6 mAb and anti-Pax5 pAb at 1:1,000 dilutions at room temperature for 1 h and rinsed twice with PBS. The cells were then stained with secondary Alexa Fluor 488 or Alexa Fluor 565-labeled monkey anti-mouse IgG antibody and Alexa Fluor 488-labeled goat anti rabbit IgG antibody at 1:1,000 dilution. Finally, the nucleus was stained by 4′,6-diamidino-2-phenylindole (DAPI) (Abcam, China). Sections were visualized under a confocal laser scanning immunofluorescence microscopy (Leica SP8, German).

### Viral load by qPCR

Absolute qPCR was conducted to determine MRV genome DNA copy using DNA templates isolated from WBC or purified B, T, macrophage and residual WBC through FastPure Cell/Tissue DNA Isolation Mini Kit (Vazyme, China) according to the manufacturer’s instructions. An absolute standard curve of MRV *mcp*-specific qPCR system was established using a 10-fold serially diluted pMD-19T-MCP recombinant plasmid DNA (10^9^copy/μL – 10^1^copy/μL). The MRV specific primer set, used in this study, targeted a partial fragment of the *mcp* gene was shown in table 1. The reactions were performed using the Light Cycler® 480-II Multiwell Plate 384 real-time detection system (Roche Diagnosis, the USA) under the following conditions: 1 cycle at 95°C for 60 s, and 40 cycles of 95°C for 5 s, 60°C for 30 s and 72°C for 5 s. A non-template control was used as a negative amplification control. All real-time qPCRs were performed in triplicate.

**Table 1.**
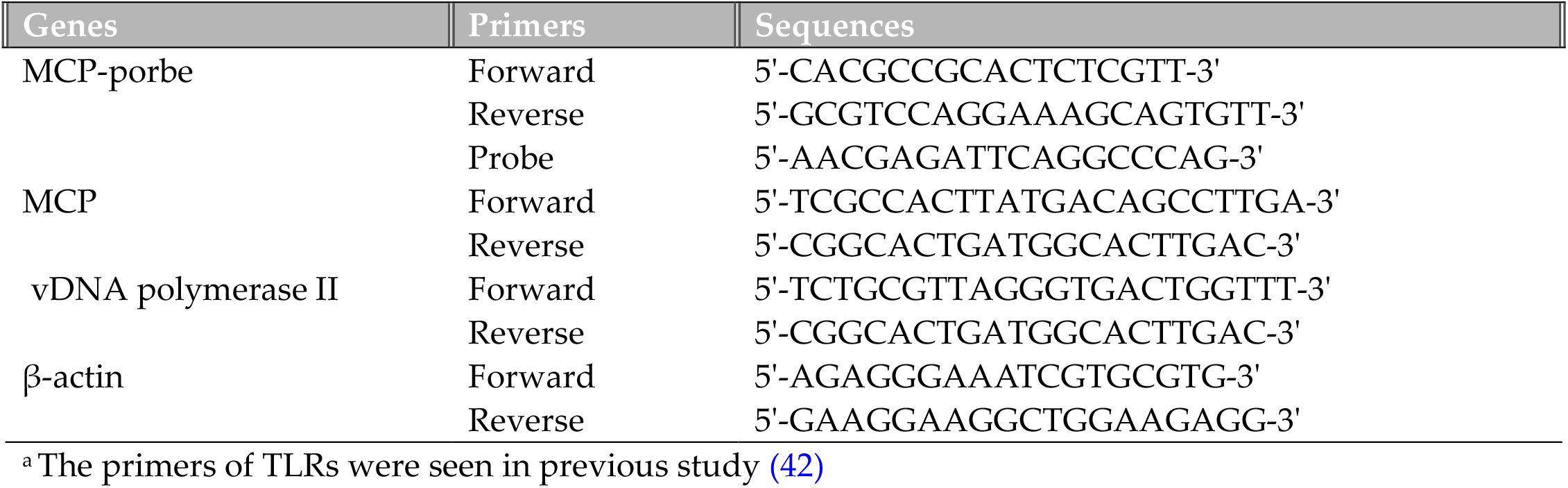
PCR prime sequence ^a^.

### Quantitative gene expression analyses

Total RNA was extracted from WBC by using Eastep Super Total RNA Extraction Kit (Promega, China) according to the manufacturer’ s instruction. Then RNA was reverse-transcribed into cDNAs using Evo M-MLV RT Premix for qPCR (Accurate Biology, China). The expressions levels of the MRV *mcp* gene and 16 different TLRs in the cDNA samples were quantified by qRT-PCR methods in the Roche LightCycler 480 system, with mandarinfish β-actin as the reference gene. Primers specific for 16 different TLRs were designed and validated by gradient PCR and with the qPCR melting curves. All primers used in the study are listed in Table 1. The qRT-PCR was performed as described previously(14). Briefly, the PCRs were performed as the following procedure: 95 °C for 30 s, one cycle; 95 °C for 5 s and 60 °C for 30 s, 40 cycles; 95 °C for 5 s, 60 °C for 1 min and 95 °C, one cycle; 50 °C for 30s, 1 cycle with a total reaction volume of 10 μL, containing 1 μL of cDNA, 5 μL of 2 × SYBR Green Pro Taq HS Premix (Accurate Biology, China), 0.5 μM primers (Tsingke, China) and 3 μL of RNase free water.

### Immune-histochemistry assay (IHC)

Tissues of moribund fish from treatment with temperature stress, vaccination or dexamethasone stimulation were collected and fixed with alcohol-formal-acetic (AFA) for immune-histochemistry assay (IHC) analysis as our previous description (14). Briefly, intestines or pyloric caecum was dissected and fixed using alcoholformalacetic and then embedded in paraffin wax. The embedded tissues were excised into sections of 4 μm thickness. The sections were dewaxed in xylene, rehydrated in a series of ethanol. Positive sections were performed using anti-MRV mAb 1C4 (1:1000) as the primary antibody and horseradish peroxidase (HRP)-labelled Goat anti-mouse IgG as the second antibody for IHC analysis. Finally, the tissue sections were developed with DAB solution and visualized under a Nikon fluorescence microscope (Eclipse Ni-E, Japan).

### Statistical analysis

Analysis of variance (ANOVA) was performed for statistical analysis of expression and viral load data. Statistical analysis of survival data was performed using a Log-Rank Test (GraphPad Prism 8, San Diego, CA, USA). A probability value of p < 0.05 was used in all analyses to indicate significance. Error bars on all graphs represent the standard error of the mean (SEM)

## RESULTS

### Transition of MRV from acute infection to persistently covert state

Two groups of mandarinfish rearing in two different temperatures were challenged with a sublethal dose (10^6.5^ TCID_50_/fish) of MRV-ZQ17. As a result, the infected fish at 26°C group occurred death at 6 dpi until no additional death occurrence at 12 dpi with an accumulated mortality of 46.67% (7/15) (**Fig. 1B**). The dead fish were confirmed as MRV infection by conventional PCR detection. By contrast, no mortality and mortality were observed at 19°C group **(Fig. 1B)**. Observation was continued to thirty days, and no additional death occurred in both groups. At 30 dpi, blood was pooled from all survival fish to isolate WBC for MRV genome DNA detection and no MRV was detected by conventional PCR, whereas low copies of MRV genome DNA were obtained from all 28 fish by absolute qPCR. The average viral loads in WBC from 20 mandarinfish rearing at 19°C was 18.5±9.7 genome equivalents (GEs) per microgram DNA (**Fig. 1C)**, whereas it was 33.9±20.7 GEs per microgram DNA from 8 survival mandarinfish at 26°C (**Fig. 1D**). Additionally, using anti-MRV mAb-based colloidal gold-immunochromatographic fast detection strip, blood sample from the survival fish showed negative result, whereas it was strong positive result for ascites sample from moribund mandarinfish in group I (**Fig. 1E**).

### Re-activation of covert MRV via raising temperature stress

Following the initial blood drawing for WBC separation at 30 dpi, mandarinfish were allowed to recover for 7 days. Subsequently, raising temperature stress was conducted in 19°C infection group. As a result, total of 5 mandarinfish (designated as M2, M8, M11, M13, and M15) died successively on the 10th, 12th, 14th, and 15th days (from 24°C to 26°C). Among them, obvious ascites was just observed in M13. Moreover, high levels of MRV loads were measured in various organs of M2, M8 and M13 (**Fig. 2B**), indicating these fish died of MRV infection. Immunohistochemistry also demonstrated a substantial number of virus-positive signals, also indicating an active MRV infection (**Fig. 2C**). After raising temperature stress, the second round of blood collection was conducted at 57 dpi and WBC from 15 surviving mandarinfish were collected (**Fig. 2A**). qPCR analysis revealed that the MRV copy number in WBC from 15 surviving mandarinfish is 139±165 GEs/mg DNA (**Fig. 2D and E**).

### Reactivation of covert MRV by vaccination and dexamethasone treatment

To examine the effect of the inactivated vaccine on the reactivation of covert MRV, 8 mandarinfish survived from 26°C challenge (group I) was selected for intraperitoneal injection of 100 μL of inactivated MRV vaccine at 38 dpi. As a result, 3 fish succumbed at 6^th^, 7^th^ and 12^th^ days post-vaccination (**Fig. 3A**). Quantitative PCR analysis confirmed significant viral load changes in all three dead fish (**Fig. 3B**), suggesting that the inactivated MRV vaccine reactivated the covert MRV to partial fatal outcomes. A repeated experiment involving 20 covert-infected individuals injected with MRV vaccine resulted in an accumulative mortality of 35% (7/20) (**Fig. 3E**). Quantification PCR of tissues from the deceased fish showed high levels of MRV loads (**Fig. 3F**). Immunohistochemistry also exhibited a substantial number of virus-positive signals in the pyloric caecum (**Fig. 3H)**.

**FIG. 3.**
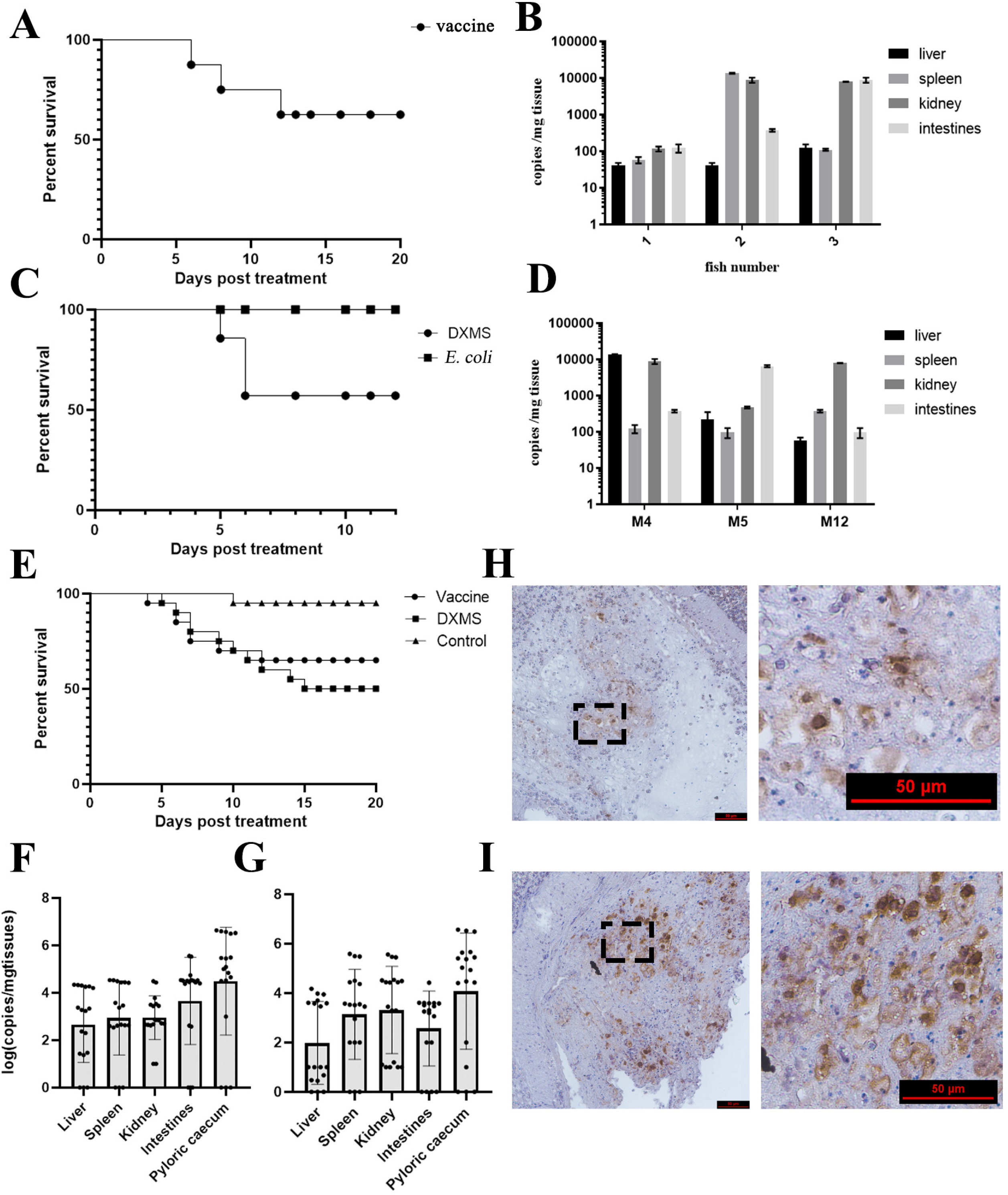
Assessment of the reactivation effects of covert MRV via vaccination, dexamethasone stimulation and *E. coil* injection. (A) Survival curve of PCI mandarinfish via vaccination stimulation. (B) Tissue MRV DNA load examination of 3 dead fish suffering vaccination stimulation. (C) Survival curves of PCI mandarinfish via dexamethasone treatment and *E. coil* injection. Three fish died during dexamethasone treatment and no fish died during *E. coil* injection. (D) Tissue MRV DNA load examination of 3 dead fish suffering dexamethasone treatment. (E), (F) and (G), The repeated experiments (N=20) of assessment of the reactivation of MRV PCI mandarinfish via vaccination and dexamethasone stimulation, and 1, 7 and 10 fish were dead in control, vaccination and dexamethasone treatment. High tissue MRV DNA loads were determined in dead fish from vaccination (F) and dexamethasone treatment (G). Strong positive signals by IHC were also observed in dead fish from vaccination and (H) and dexamethasone treatment (I).

At 64 dpi, 14 of surviving mandarinfish in group II were divided into two groups, 7 fish were received intraperitoneal injection of 100 μL of dexamethasone and another 7 fish were intraperitoneally injected with heat-inactivated *E. coli*. Daily observation revealed that one and two fish died at 5th day and 6th day post dexamethasone treatment (M4, M5 and M12), respectively (**Fig. 3C**). Quantitative PCR analysis confirmed high viral loads in these dead fish (**Fig. 3D**). A repeated experiment with 20 covert MRV mandarinfish treated with dexamethasone resulted in an accumulative mortality of 50% (10/20) (**Fig. 3E**). qPCR quantification of tissues from deceased fish showed high levels of MRV genome DNA (**Fig. 3G**), and immunohistochemistry also displayed a significant number of virus-positive signals in the pyloric caecum (**Fig. 3I**). All these results suggested dexamethasone’s ability to reactivate covert MRV to fatal outcomes. No dead fish was observed in *E. coli* injection group (**Fig. 3C**). Blood samples were collected from all survival fish at 7th day post *E. coli* treatment for conventional PCR or colloidal gold-immunochromatographic fast detection and both showed MRV negativity (data not shown), indicating that *E. coli* treatment could not activate covert MRV to a detectable level.

### Purification of B cells, T cells and *MΦ*

Flow cytometry was employed to screen and concentrate B cells, T cells, and macrophages in peripheral blood. The sorting markers included IgM and the transcription factor Pax5 (an additional marker specific to B cells) for B cells, CD3Δ for T cells, and MRC1 for macrophages. The validity of these cell markers was firstly conducted through Western blotting. The result showed that all four antibodies could recognize these markers from WBC on their expected positions **(Fig. 4A)**. The subsequent flow cytometry analyses demonstrated that in peripheral blood WBC of covert MRV-carried fish, IgM^+^ B lymphocytes constituted approximately 25±7.3%, CD3Δ^+^ T lymphocytes accounted for around 24±4.6%, and MRC1^+^ macrophages made up about 5±2.4% (**Fig. 5A**). Following sorting by flow cytometry, the percentage of IgM^+^ B lymphocytes increased from 31.4% to 83.4%, CD3Δ^+^ T lymphocytes increased from 22.1% to 93.7%, and MRC1^+^ macrophages increased from 5.1% to 88.3% (**Fig. 3E-G**). The concentration process of B cell was assessed by anti-IgM mAb-based and anti-Pax pAb-based immunofluorescence assay (IFA) by using confocal microscopy. As shown in **Fig. 4B**, only a few WBC could be stained by anti-IgM mAb in native WBC (**Fig. 4B1**), however, the majority of cells were recognized by anti-IgM mAb after FACS sorting (**Fig. 4B2 and C**). By contrast, almost no fluorescence signal was observed in the residual WBC out of the isolation of B cells (**Fig. 4B3**). All these results indicated B cells were well isolated and purified from WBC. Additionally, the co-expression of sIgM and Pax in the purified B cells were also observed (**Fig. 5B**). In agreement with FACS analysis, most flow cytometry-sorted IgM^+^ WBC exhibited IgM staining on the cell surface, whereas Pax5 staining localized in the nucleus (**Fig. 4B and C; Fig. 5B**).

**FIG. 4.**
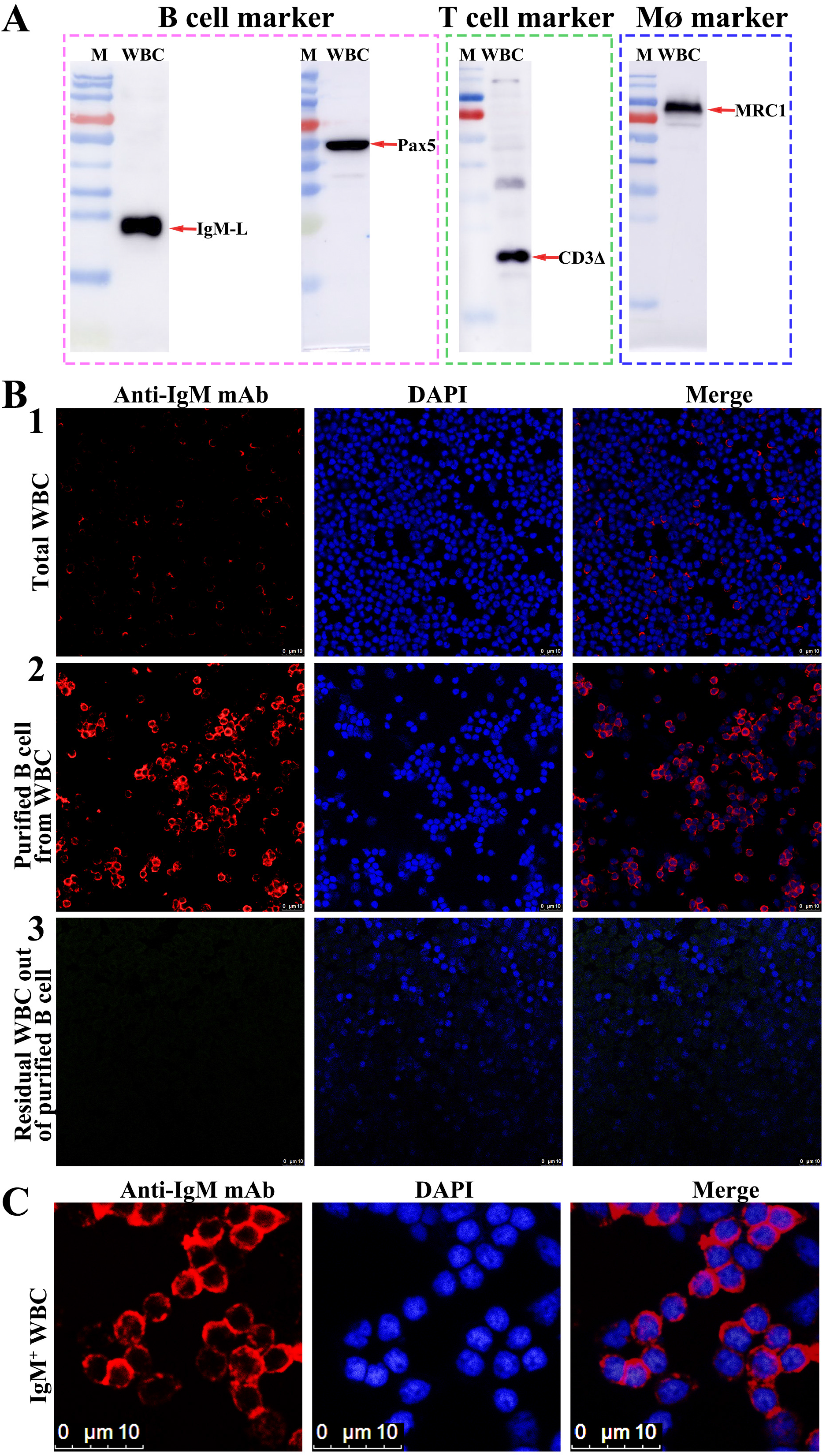
Detection of the antibody-targeted cell markers from mandarinfish WBC. (A) Western blotting analysis of four cell markers of mandarinfish WBC. The mAb of 7F12F6 recognize the light chain of mandarinfish IgM. Three pAbs also recognized the corresponding protein bands as speculated; (B) and (C) Confocal micrographs of mandarinfish WBC. (B1), a few IgM^+^-labeled B cells from native WBC; (B2), numerous IgM^+^-labeled B cells after flow cytometry sorting; (B3), almost no IgM^+^-labeled B cells in the residual WBC out of B cells. (C) The enlarged image of sorted IgM^+^-labeled B cells by FACS.

**FIG. 5.**
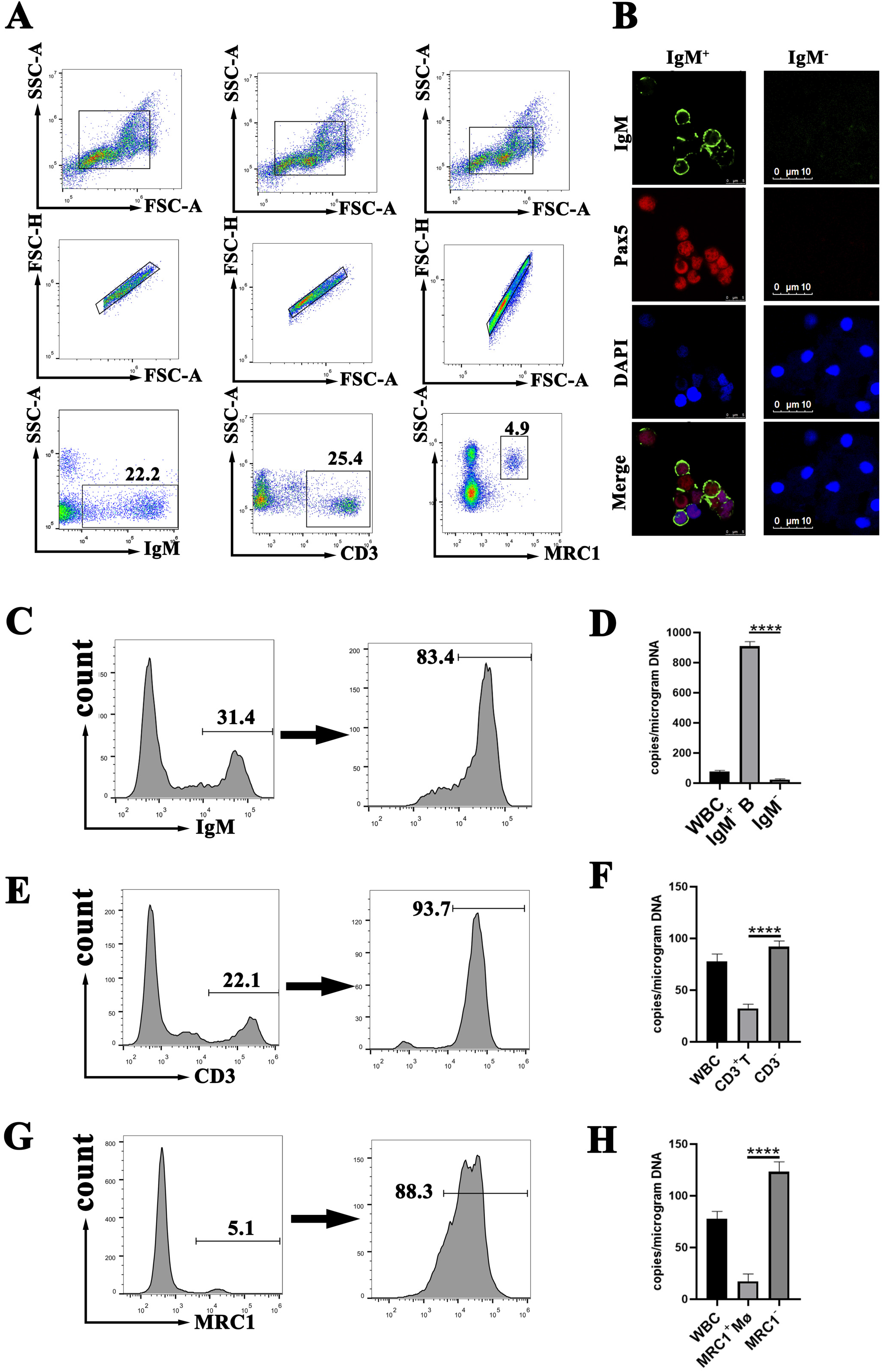
Fluorescence-activated cell staining (FACS) sorting of mandarinfish peripheral WBC. (A) Flow cytometry of mandarinfish WBC stained with antibodies to mandarinfish IgM (left), CD3 (middle), and MRC1(right). (B) Confocal micrographs of mandarinfish IgM^+^ and IgM^-^ WBC. IgM^+^ cells were identified by anti-mandarinfish IgM mAb (7F12F6) and secondary Alexa Fluor555-labelled goat anti-mouse IgG (green). Pax5^+^ cells were identified by anti-Pax5 pAb and secondary Alexa Fluor488-labelled goat anti-rabbit IgG (red). The nucleus was identified with DAPI (blue). The merged images show cells visualized by all three fluorescence. (C) and (D) indicate the progress of IgM^+^ WBC selected by FACS and the copy number of MRV genomes determined from 1 μg total DNA of total WBC, the selected IgM^+^ B cells and the residual IgM^-^ WBC from MRV PCI mandarinfish, respectively; (D). (E) and (F) indicate the progress of CD3^+^ WBC selected by FACS, and the copy number of MRV genomes detected from 1 μg total DNA of total WBC, selected CD3^+^ T cells and CD3^-^ WBC from MRV PCI mandarinfish, respectively. (G) and (H) indicate MRC1^+^ WBC selected by FACS (G) and the copy number of MRV genomes detected from 1 μg total DNA of total WBC, selected MRC1^+^ Mø and MRC1^-^ WBC from MRV PCI mandarinfish, respectively.

### MRV genome assessment in B cells, T cells and macrophages

To determine which WBC type is the targeted cell response for the covert MRV, the MRV genome copies in purified B cells, T cells, and macrophages from PCI mandarinfish were determined by real-time qPCR. As a result, under an equal amount of total template DNA, the MRV genome DNA loads in purified IgM^+^ B cells was about 40-fold abundant than those in IgM^-^ WBC (**Fig. 5D**). By contrast, the MRV genome loads in CD3Δ^+^ T cells and in MRC1^+^ macrophages were only approximately one-third and one-tenth of those in CD3Δ^-^ WBC and MRC1^-^ WBC, respectively (**Fig. 5F and H**), clearly indicating that B lymphocyte but not T lymphocyte and macrophage was the targeted cell for the establishment of covert MRV.

### Screening for TLR pathway involvement in the reactivation of covert MRV

To investigate the possible involvement of particular TLR pathway upon the reactivation of covert MRV, the differential expression of TLR genes during the transition of covert MRV to reactivation was assessed.

Mandarinfish (weighing ∼250g) was infected with a sublethal dose (**10**^**6.5**^ TCID_50_/fish) of MRV and maintained for 30 days to ensure the establishment of PCI as the above description. At 30 dpi, the PCI mandarinfish were injected with dexamethasone as the above-mentioned approach. The injection of equal PBS was set as a control. The blood was pooled and the peripheral WBC was isolated at 1 day and 3 days post treatment. Absolute qPCR was conducted to determine the viral genomic load. Relative quantification PCR was conducted to determine the transcription expression of viral MCP and DNA polymerase as well as 16 different TLRs. As a result, the MRV genomic load in WBC increased significantly (n < 0.1) regardless of dexamethasone treatment at 1 day and 3 days **(Fig. 6A)**. At transcription level, significant upregulation expression of viral MCP and DNA polymerase were also measured **(Fig. 6B and C)**, indicating the covert MRV was reactivated to replicate upon dexamethasone treatment. Meanwhile, all TLRs except for TLR2A showed inductive effects with the expression level increasing by less than 1.5 times (**Figure 6D**).

**FIG. 6.**
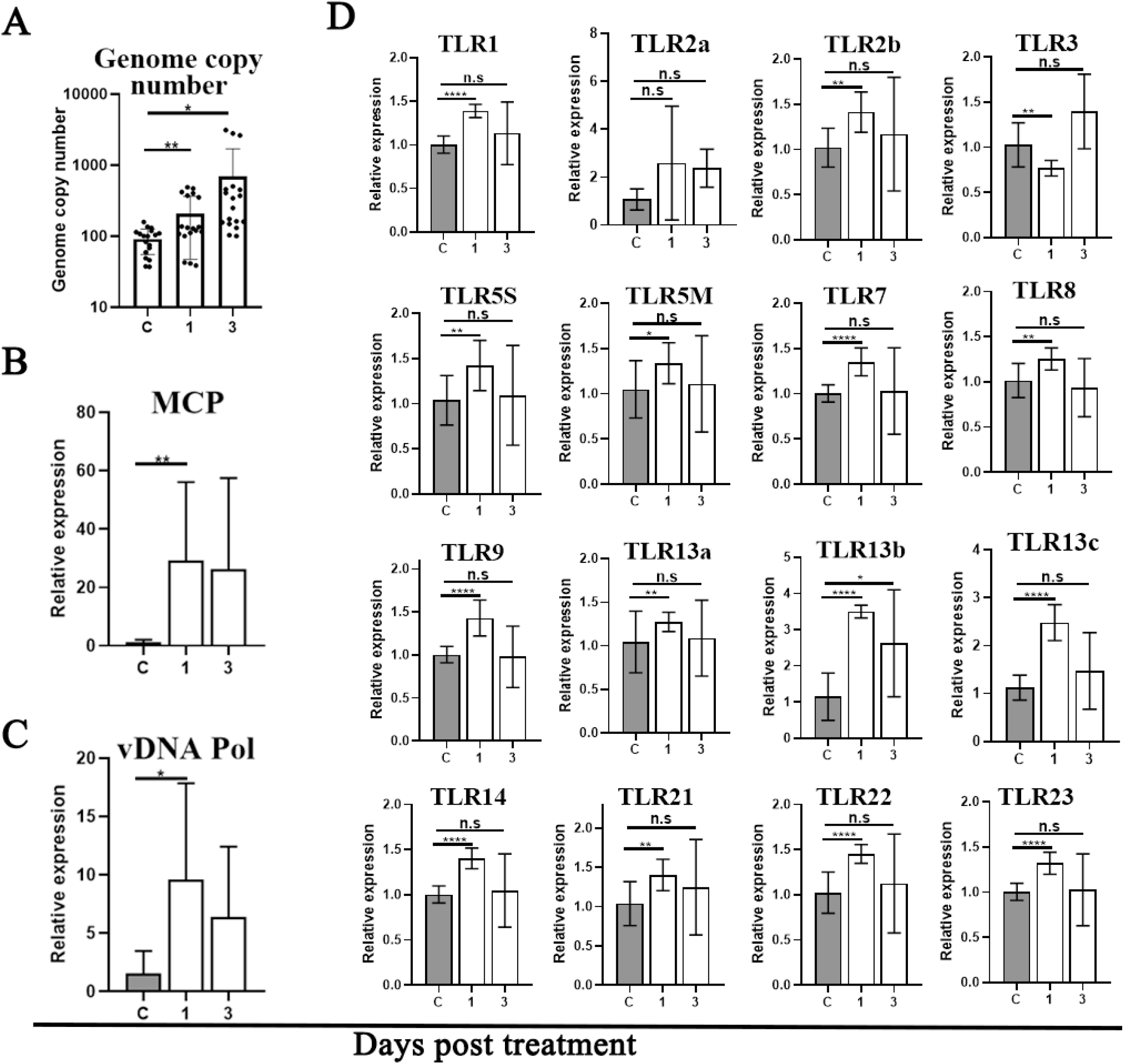
Differential expressions of TLR genes in WBC of MRV PCI mandarinfish following dexamethasone stimulation. (A) MRV genome DNA copy number of WBC in control, 1 day post treatment and 3 days post treatment. (B) and (C) Relative *mcp* gene and DNA polymerase gene expression in control, 1 day post treatment and 3 days post treatment, respectively. (D) Relative gene expression levels of 16 TLRs of WBC in control, 1 day post treatment and 3 days post treatment. Statistical significance between control and treated groups is denoted by *, where the P value was,0.05 using one-way ANOVA and Tukey’s post hoc test. ND, not determined.

Given that *E. coli* and flagellate-stimulated TLR5 are implicated in the activation of quiescent FV3 infection(38), the homologous ligands were further used to determine the stimulation specificity of TLR5 in mandarinfish WBC. As a result, intraperitoneal injection of both inactivated *E. coli* and flagellin led to a sharp upregulation expression of TLR5M transcriptions in WBC at 1 and 3 dpi (**Fig. 7B and E**).

**FIG. 7.**
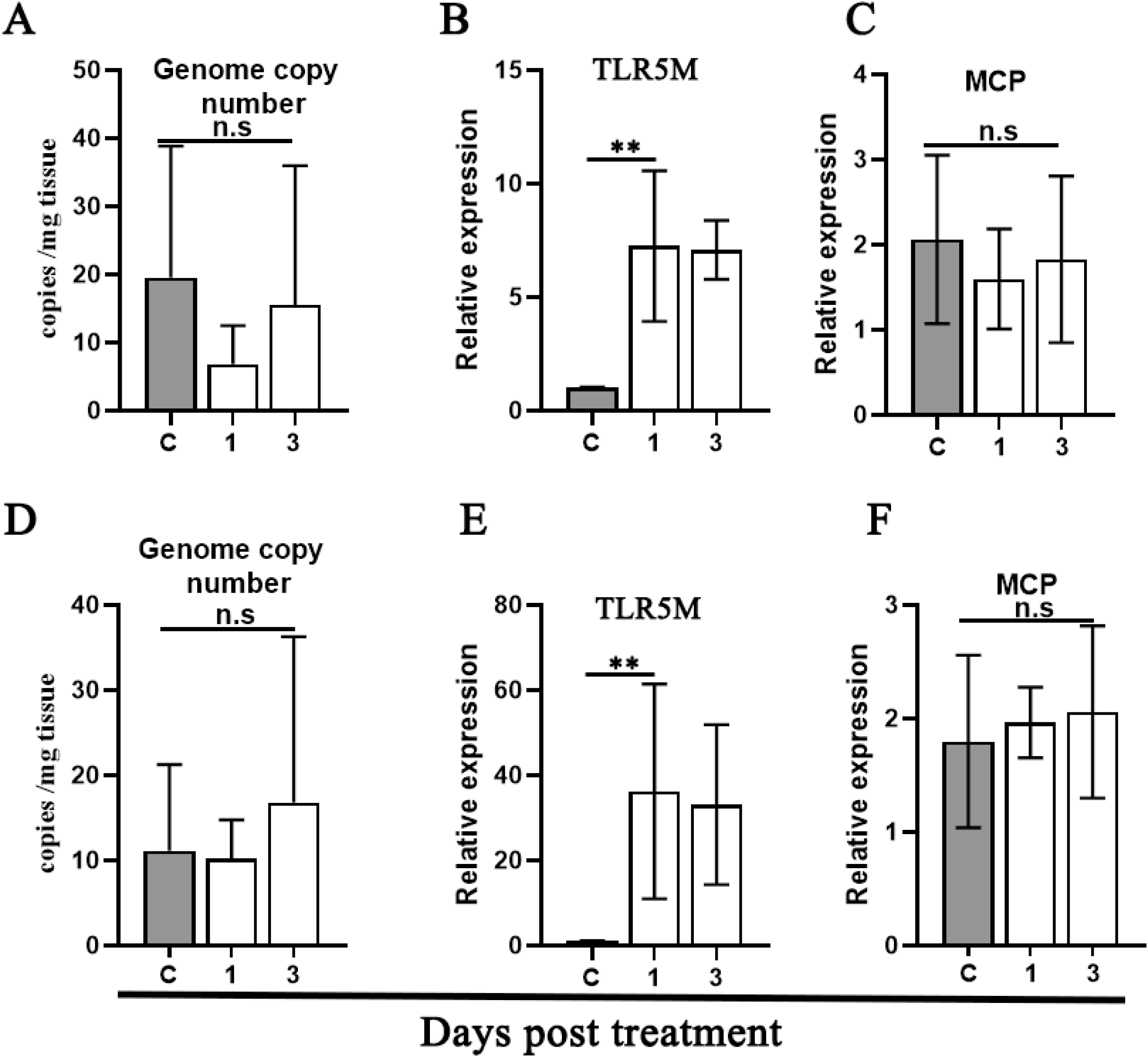
Assessment of the reactivation of covert MRV and TLR5 signal by stimulation with *E. coli* and flagellin. MRV genome copy number(A), Relative TLR5M gene expression (B) and relative MCP expression (C) upon *E. coli* stimulation. MRV genome copy number (D), Relative TLR5M gene expression (E) and relative MCP expression (F) upon flagellin stimulation. Relative gene expression levels of WBC from MRV covert fish stimulation with *E. coli* and flagellin for 1 or 3 days or injected with PBS as a control. Statistical significance between control and treated groups is denoted by *, where the P value was,0.05 using one-way ANOVA and Tukey’s post hoc test. ND, not determined.

However, no increase of MRV genome DNA copy and no upregulation expression of *mcp* RNA were determined (**Fig. 7A-F)**, indicating that *E. coli* and flagellate stimulation really initiated TLR5 pathway as expected, however, the active TLR5 was not linked to the reactivation of covert MRV.

## DISCUSSION

The natural persistent infections of MRV have been investigated and characterized in our previous report(14). The core points of persistent MRV infection includes: (a) it is a long-term chronic infection with a disease duration up to 5 months, almost covering the entire breeding cycle of commercial mandarinfish, (b) during persistent infection, the MRV load in various visceral organs is too low to be detected by conventional PCR, and (c) the featured clinical signs are lethargy, anorexia and body emaciation and deformity accompanied with sporadic low mortality. Furthermore, our recent study showed that the o/w formation of inactivated MRV vaccine conferred no protection against the parental virus challenge, which were also confirmed by several other research teams (data not shown). Thus, the underlying mechanism of persistently covert infection and the ineffectiveness of inactivated vaccines have become the major issues currently troubling the understanding of MRV.

For cold-blooded animals, many habitat characteristics such as water chemistry, soil type, ambient temperature, hydroperiod, UV-B as well as population density can affect the onset of viral diseases (7). In artificial challenge of ISKNV, the water temperature of 20°C was experimentally evidenced as the threshold temperature(43). Over 20°C, ISKNV showed highly lethal to mandarinfish regardless of various infection routines including administration by intraperitoneal injection, intramuscular injection or cohabit manner, however, ISKNV never showed lethality to mandarinfish when the experimental temperature was below 20°C. In this study, our data showed that 19°C is also a nonpermissive temperature for the onset of MRV disease, whereas 26°C is a permissive temperature for MRV outbreak. Under a sublethal dosage of MRV challenge, no fish died (0/20) in 19°C group, but 46.7% (7/15) fish died of MRV infection accompanied with ascites symptoms at 5 dpi in 26°C group (**Fig. 1B**). The MRV could be easily detected through colloidal gold-immunochromatographic fast detection strip (**Fig. 1E**), conventional PCR as well as MFF-1-based virus isolation. At 30 dpi, the average viral loads in WBC of 20 mandarinfish rearing at 19°C was 18.5±9.7 genome equivalents (GEs) per microgram DNA (**Fig. 1C)**, whereas it was 33.9±20.7 GEs per microgram DNA from 8 survival mandarinfish at 26°C (**Fig. 1D**). Since no additional mortality occurred and the viral load in blood is too low to be detected by conventional PCR and colloidal gold-immunochromatographic fast detection strip, it is proposed all these fish entered covert infection phase. Furthermore, the result also suggested that MRV infection under nonpermissive temperature of 19°C could spontaneously entry into persistent covert infection state. By contrast, under permissive temperature of 26°C, infection with a sub-lethal dosage of MRV could result in partial mortality, and the survival then transit into persistent state.

During the gradually raising temperature, 5 fish died, and 3 of them were measured to have detectable high viral load (**Fig. 2B**). Absolute qPCR showed that the average viral load (N=15) in WBC from the survivals was 139±165 GEs/mg DNA (**Fig. 2E**), which is much higher than that (18.5±9.7 genome equivalents (GEs) per microgram DNA) from fish (N=20) before temperature stress (**Fig. 2D and E**). The result again indicated that the raising temperature stress could reactivate covert MRV to replicate and result in fatal outcome and then the survival once again entered persistent state of infection. After two rounds of drawing blood and following another 7 day’s stabilization, the survival fish were divided into two groups and treated with dexamethasone and heat-killed *E. coil*, respectively (**Fig. 1A**). As a result (**Fig. 3C**), mortality was observed in dexamethasone-treated group but not in heat-killed *E. coil-* treated group. Both viral load determination and IHC analysis demonstrated that dexamethasone functions as an effective inducer to trigger the reactivation of persistent MRV, which were confirmed by two independent repeated experiments in this study (**Fig. 3E and Fig. 6**). Dexamethasone, a classic glucocorticoid immunosuppressants, was commonly used in the reactivation of various viruses with persistent or latent characteristics (44-46). For instance, cyprinid herpesvirus 2 (CyHV-2), a highly contagious pathogen of goldfish (*Carassius auratus*) and Prussian carp (*Carassius auratus gibelio*), establishes persistent infection in goldfish monocytes/macrophages. Dexamethasone could induce the persistent CyHV-2 to replicate via effectively suppressed monocyte/macrophage function and antibody production(44). The mechanism of dexamethasone-induced reactivation of covert MRV remained unclear and would be investigated in future.

Previous study showed that vaccination with the o/w formation of inactivated ISKNV vaccine could induce mandarinfish with covert MRV to mass mortality(14). This study showed that the o/w formation of inactivated MRV vaccine also induced similar outcome for covert MRV. It was proposed that the white oil component on one hand plays adjuvant role in w/o formation of emulsified vaccine, and on the other hand, the obvious side effect of o/w formation of vaccine is the induced local fibroperitonitis. The induced peritonitis might trigger the reactivation of covert MRV. Generally, this study once again demonstrates the presence of a persistently covert infection of MRV in mandarinfish. Furthermore, the covert MRV can be triggered reactivation through various pathways, encompassing temperature stress, vaccination, and the immunosuppressive agent dexamethasone stimulation.

Numerous epidemiological investigations and clinical data suggested that ranavirus has the capability to establish asymptomatic infections and reactivation under specific conditions(7, 32, 36, 37), however, the exact persistent infection has only been validated in FV3. In case of FV3, the peritoneal macrophages was confirmed the major reservoir for quiescent FV3, and the quiescent FV3 can be reactivated by *E. coli* flagella-mediated TLR5 signal pathway(36, 38). In mammalian systems, human herpes simplex virus (HSV) establishes latency in ganglia, reactivating under reduced body resistance or various stress-induced host immune states(25). Similarly, Bovine herpesvirus 1 (BoHV-1) achieves latency in the spleen, tonsils, and groin lymph nodes, reactivating after various stress stimulation (45). While γ-herpesviruses, including human Epstein-Barr virus (EBV) and murine γ-herpesvirus 68 (MHV-68), establish latent infection in B lymphocytes (47). In fish, CyHV3/KHV establishes latency in B lymphocytes of koi carp, reactivating in response to temperature stimulation and physical stress(26, 48), whereas CyHV2/GFHNV establishes persistent infection in goldfish monocytes/macrophages and could be reactivated by dexamethasone(44). In this study, we employed specific antibodies against MRC1 to isolate macrophages and purified T cells and B cells by using specific antibodies against CD3 and IgM. MRC1, a pattern recognition receptor, serves as a marker for macrophages and has been identified in various fish species(49, 50). CD3, exclusive to T cells, is composed of 6 peptide chains and participates in T cell antigen recognition and signal transduction(51). IgM, the principal component of fish immunoglobulins produced by B cells, functions as an antigen receptor on B lymphocytes. To determine the identity of the selected IgM^+^ cells, an antibody specific to B cell transcription factors Pax5 was also used in our study. The transcription factor Pax5 is a major regulator of B cell development and has been identified in both mammalian and non-mammalian species (52, 53). Pax5 is mainly expressed in vertebrate B-cell lineages(54), including bony fish species such as rainbow trout, puffer fish, and zebrafish(55-57). In our study, the whole blood cell from PCI mandarinfish was sampled for MRV load determination and WBCs but not red blood cells were determined as the targeted cells. Furthermore, using these anti-mandarinfish marker antibodies, B cells, T cells and macrophage were well isolated and purified from WBC (**Fig. 5A and B**). qPCR showed that MRV genomic DNA load in IgM^+^ B cells was 40 times higher than that in IgM^-^ WBC after isolation. By contrast, the MRV genomic loads in CD3^+^ T cells and MRC1^+^ macrophages were much lower than those in WBC and negative cells (**Fig. 5F**). All these results showed that peripheral blood IgM^+^ B lymphocytes serves as the primary target cells for the covert infection of MRV.

To assess the possible TLR signals involving the reactivation of covert MRV, 16 mandarinfish TLR molecules were screened. Under dexamethasone treatment, the covert MRV was reactivated as expected with slight upregulation expression of 15 of 16 TLR genes (**Fig. 6**). Meanwhile, when the PCI mandarinfish was injected with heat-killed *E. coli* or flagella, the TLR5 signal was significantly activated as scheduled, however, the covert MRV was not reactivated (**Fig. 7**). All these results suggest that the reactivation of covert MRV is a non-TLR5 manner, which is much different from that of persistent FV3 (38). The dissimilarity in their activation pathways may be attributed to their distinct taxonomic status and host preferences, despite both belonging to the *Ranavirus* genus within the *Iridoviridae* family. Specifically, ranaviruses can be classified into two subclasses: FV3 cluster in the ALRV subclass, primarily infecting amphibians and reptiles, and MRV cluster in LMBV-Like, which alongside GIV-like, only infects bony fish(5). At genomic level, the ALRV subclass genome is approximately 105 kb(1, 58), the GIV genome is about 140 kb(59), and the MRV genome is only about 98kb(3, 4). Notably, the homology between MRV and ALRV main capsid protein is only 84%. These genomic differences maybe contribute to the heterogeneity observed in fish ranavirus infections compared to amphibian ranavirus infections(3).

It is worth noting that the preexposure to virulent megalocytiviral ISKNV/RSIV in nonpermissive temperature could elicit robust protective immunity against the re-challenges with parental viruses in permissive temperature, thus so-called live ISKNV/RSIV vaccines were developed on the basis of temperature regulation(22, 60). As for MRV, the low copy of MRV genome DNA in nonpermissive temperature of 19°C at least maintained for 30 days. Even if a mild warming stimulation was applied, partial mortality outcome occurred and the survival again enter persistent status. Finally, the covert MRV could be once again reactivated by dexamethasone treatment (**Fig. 3C**). Obviously, the exposure to virulent live MRV under nonpermissive temperature could not induce enough immune clearance, which is high consistent with field observation. In production practice, the surviving fish from the ISKNV epidemic usually do not experience a recurrence of ISKNV infection, however, for MRV, after a primary infection, the surviving fish usually maintains a chronic infection state, which can last for at least 5 months(14). Is B cell target persistent infection a key factor affecting activated immune clearance worth further investigation. Specifically, in contrast to the highly effective inactivated ISKNV vaccine, robust experimental evidences show that the o/w formation of inactivated MRV vaccine confers no protection against the challenge with virulent MRV regardless of antigen dosage. Whether the complete ineffectiveness of inactivated vaccines is associated with the persistent targeting of MRV specific B cells and its underlying mechanisms will be an important topic in the study of vaccine prevention of MRV and its pathogenesis.

## ACKNOWLEDGMENTS

This work was funded by Guangdong Basic and Applied Basic Research Foundation under No. 2023A1515010009; The key areas R&D Program of Guangdong Province under No. 2021B0202040002; Innovation Group Project of Southern Marine Science and Engineering Guangdong Laboratory (Zhuhai) under No. 311021006 and Guangdon g Provincial Special Fund for MAITIT [Grant Number 2019KJ141].

